# Cysteine-Rich Intestinal Protein 1 is a Novel Surface Marker for Myometrial Stem/Progenitor Cells

**DOI:** 10.1101/2023.02.20.529273

**Authors:** Emmanuel N. Paul, Tyler J. Carpenter, Sarah Fitch, Rachael Sheridan, Kin H. Lau, Ripla Arora, Jose M. Teixeira

**Author notes:** **Corresponding Author:** Jose M. Teixeira, PhD, Michigan State University, 400 Monroe Ave NW, Grand Rapids, MI 49503. Acknowledgments: The authors have no conflicts to declare.

## Abstract

Myometrial stem/progenitor cells (MyoSPCs) have been proposed as the cells of origin for uterine fibroids, which are benign tumors that develop in the myometrium of most reproductive age women, but the identity of the MyoSPC has not been well established. We previously identified SUSD2 as a possible MyoSPC marker, but the relatively poor enrichment in stem cell characteristics of SUSD2+ over SUSD2- cells compelled us to find better discerning markers for more rigorous downstream analyses. We combined bulk RNA-seq of SUSD2+/- cells with single cell RNA-seq to identify markers capable of further enriching for MyoSPCs. We observed seven distinct cell clusters within the myometrium, with the vascular myocyte cluster most highly enriched for MyoSPC characteristics and markers, including *SUSD2. CRIP1* expression was found highly upregulated in both techniques and was used as a marker to sort CRIP1+/PECAM1- cells that were both enriched for colony forming potential and able to differentiate into mesenchymal lineages, suggesting that CRIP1+/PECAM1- cells could be used to better study the etiology of uterine fibroids.

## Introduction

Uterine fibroids, also known as leiomyomas, are benign tumors found in the smooth muscle layer of the uterus, the myometrium. Uterine fibroids develop in up to 80% of women during their reproductive years and, although benign, are often associated with debilitating symptoms such as menorrhagia, anemia, dysmenorrhea, pelvic pain, and urinary incontinence (1,2). Hormonal therapies, mainly used for alleviating fibroid symptoms, are generally short-term treatments due to long-term side effects or induced infertility (3). Hysterectomy, the most common and effective treatment for uterine fibroids, results in permanent infertility (4). Progress in the search for effective medical therapies that preserve fertility and avoid invasive surgery has been difficult, in part because fibroid etiology and pathogenesis of the disease is unclear.

A dysregulated myometrial stem/progenitor cell (MyoSPC) has been proposed as the cell of origin for uterine fibroids. After embryonic development, tissue-specific stem cells remain throughout the body and play important roles in tissue homeostasis, including replacing dying cells and participating in tissue remodeling (5). The dramatic remodeling that occurs during pregnancy and following parturition in the uterus suggest a need for and the existence of myometrial stem cells (6). Uterine fibroids are thought to be a clonal disease (7–9), and since most clonal diseases have a single cell origin (10), we and others (11,12) have hypothesized that a mutated MyoSPC could be the cell of origin for uterine fibroids (13). Thus, the identification of the MyoSPC has been an important goal of many laboratories to begin studying the underlying mechanisms of fibroid etiology. The presence of cells with stem cell properties has been demonstrated using the label-retaining cells in a mouse model (14,15) and using the side population (SP) method in human myometrium (16,17). Putative MyoSPCs have been isolated and studied by using a combination of cell surface markers, including SUSD2 (18), CD44/Stro-1 (11) and CD34/CD49f/b (19). However, cell surface markers in these studies have been selected using stem cell markers from other tissues, and their respective contributions to myometrial smooth muscle regeneration have not been well established. We and others have also used the side population (SP) discrimination assay (16,17), but this stem cell identification technique has multiple pitfalls, not the least of which is difficulty in enriching and recovering live SP cells for further analyses (20). Because the endometrial stroma and the myometrium originate from the same embryonic tissue, the Müllerian duct mesenchyme (6), we recently proposed that SUSD2, an endometrium stem cell marker (21), also enriches for MyoSPCs (18). While SUSD2+ cells do have mesenchymal stem cell characteristics, SUSD2+ cells represent between 25-40% of total myometrial cells. Additionally, colony formation is only increased 2.8-fold increase in SUSD2^+^ cells compared to the rest of the myometrial cells, suggesting that further enrichment might be possible. The objective of the present study was to integrate next-generation sequencing, including single cell RNA-seq and bulk RNA-seq, to identify a more specific marker to significantly enrich for MyoSPCs from human myometrium, which can then be used to better understand the molecular mechanisms underlying fibroid etiology.

## Results

### SUSD2^+^ are enriched for characteristic MSC genes compared with SUSD2^-^ cells

To determine how best to enrich for stem cell activities in the SUSD2+ MyoSPC population, we used SUSD2 to enrich for myometrium stem cells followed by RNA-seq to discover new cell surface markers for MyoSPCs in the human myometrium. Myometrial cells from non-fibroid patients (n = 5) were isolated and live SUSD2+ and SUSD2- cells were sorted by flow cytometry. As with our previous results (18), 30-50% of the myometrial cells were SUSD2+ (**Fig 1A**). Total RNA was isolated from the two cell populations and sequenced for differential gene expression analyses. Principal Component Analysis (PCA) plot showed that SUSD2+ and SUSD2- cells were separated by principal component 1 with a variance of 39%, indicating a strong divergence in the transcriptomic profiles of these two cell populations (**Fig 1B**). A total of 6777 significant differentially expressed genes (DEGs) were detected between SUSD2+ and SUSD2- myometrial cells with a p-adjusted false discovery rate (FDR) <0.05 (**Fig 1C and Table S1**). 3527 genes were down-regulated and 3250 were up-regulated in the SUSD2+ population compared to the SUSD2- population. We confirmed that *SUSD2* was up-regulated in the SUSD2+ sorted cells and that they were also enriched in other MSC markers such as *MCAM, PDGFRβ* and *CSPG4* (**Fig 1C and 1D**). A heatmap of the top 300 DEGs from the SUSD2+ to SUSD2- cells comparison showed a good separation between cell types and included *SUSD2, MCAM, PDGFRβ* and *CSPG4* (**Fig 1E**).

**Figure 1.**
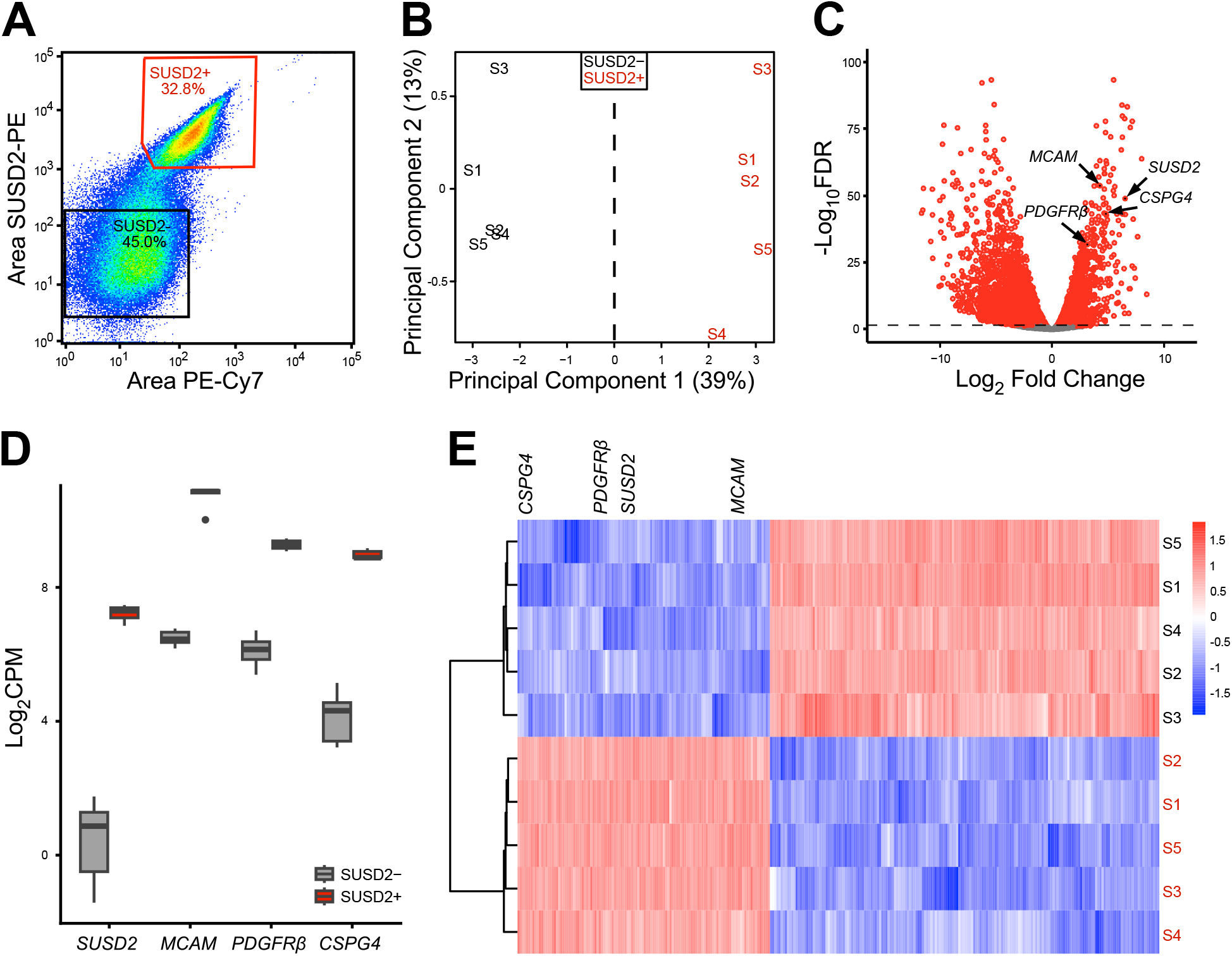
Bulk RNA-seq results of SUSD2^-^ and SUSD2^+^ cells from myometrial samples. (**A**) Representative flow cytometry scatter plot of single cell SUSD2- and SUSD2+ sort showing over 30% of the live myometrial cells positive for the SUSD2 marker. Boxed areas indicate gating strategy. (**B**) Principal component analysis (PCA) plot of RNA-seq results from SUSD2- (in black) and SUSD2+ (in red) cells, labels represent individual patient samples (n = 5), and variance for each PC is indicated in percentage. (**C**) Volcano plot showing up (n = 3527) and down (n = 3250) DEGs with an FDR-adjusted p-value < 0.05 in SUSD2+ vs. SUSD2- cells depicted as red dots. Grey dots represent genes with an FDR p-value > 0.05. (**D**) Boxplot of mesenchymal stem cells markers, *SUSD2, MCAM, PDGFRβ* and *CSPG4* in the SUSD2- (in grey) and SUSD2+ (in red) cell population (n = 5). Gene expression is shown as log_2_CPM. SUSD2+ sorted cells are significantly enriched for MSC markers including *SUSD2* (log_2_FC= 4.5, FDR p= 2.9×10^-5^), *MCAM* (log_2_FC= 3.1, FDR p= 8.8×10^-16^), *PDGFRβ* (log_2_FC= 2.8, FDR p= 1.5×10^-11^) and *CSPG4* (log_2_FC= 3.4, FDR p= 1.1×10^-11^). (**E**) Heatmap of the top 300 DEGs from SUSD2+ vs. SUSD2- cells comparison with unsupervised hierarchical clustering of genes and samples (n = 5). Color gradient represents gene expression as z-score.

### Myometrium side population cells are not enriched in MSC markers

The side population (SP) phenotype is another often used method to isolate cells with stem cell characteristics that exploits the ability of some stem cells to efflux the DNA-binding dye Hoechst 33342 via the ATP-binding cassette (ABC) transporters (11,16,17,22,23). An average of 1.7% of the total myometrial cells were SP+ (**Fig 2A**). Addition of verapamil, a calcium channel blocker used as a negative control to validate the SP, severely decreased of the number of the myometrium SP+ cells (**Fig 2B**). SP+ and SP- myometrial cells were sorted for total RNA sequencing and analyzed by PCA plot, which showed that matched SP+ and SP- cells segregated by the principal component 2, accounting for 25% of the variance (**Fig 2C**). A total of 828 significant (FDR <0.05) DEGs, including 478 upregulated genes and 350 downregulated genes, were detected between the SP+ and SP- myometrial cells (**Table S2**). The top 10 DEGs enriched in the SP+ to SP- comparison were associated with immune response (*XCL2 (24), CD69 (25), IL7R (26), KLRD1 (27*), and *IL18R1 (28)*) apoptosis (*TNFRSF10A (29)*), extracellular matrix (*SPOCK2 (30)*) and hematopoietic stem cell (*SELE (31), GATA3 (32), CD69 (33*), and *VCAM1(34)*) (**Fig. 2D**). We confirmed an increase in expression of two major ABC transporters, including *ABCB1*, and *ABCG2*, and a decrease in *PGR*, another marker previously shown downregulated in the SP+ compared to the SP- of human myometrial cells (16) (**Fig 2E**). Surprisingly, SP+ cells did not show increased expression of putative MSC markers (19,35,36), *SUSD2, MCAM, PDGFRβ, CSPG4, CD44, CD34* and *ITGA6* (also known as *CD49f*) compared to the SP- (**Fig 2F**).

**Figure 2.**
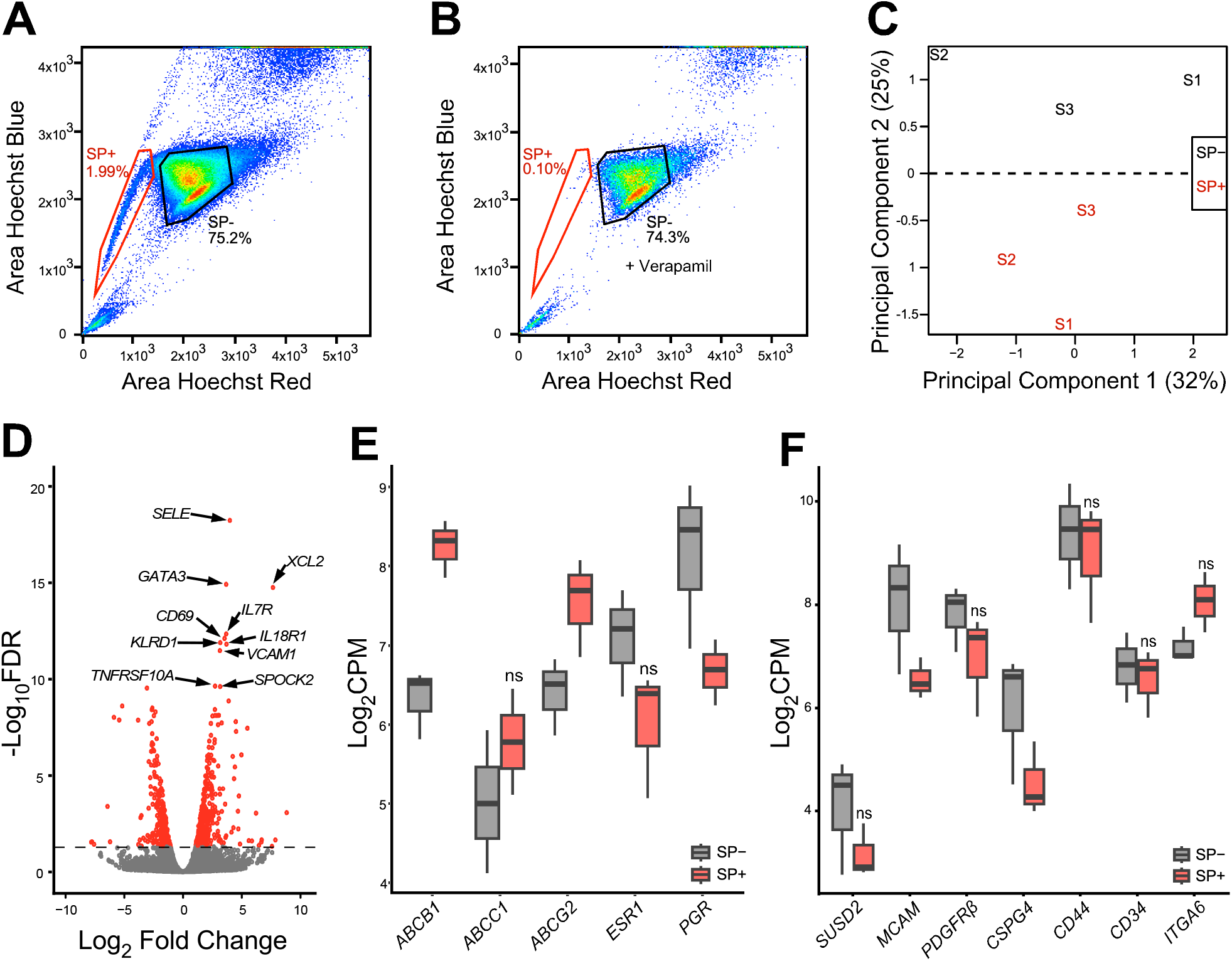
Transcriptomic analysis of the myometrium side population (SP). (**A**) Scatter plot of the gating strategy to sort the SP+ and the SP- cells from human myometrium. (**B**) Verapamil pre-treatment of myometrial cells reduces the number of the SP^+^ cells from 1.99% to 0.1% of the total live single cells. (**C**) PCA plot of RNA-seq results from SP- (in black) and SP+ (in red) cells, each label represents one sample (n = 3), variance for each PC is indicated in percentage. (**D**) Volcano plot showing up (n = 478) and down (n = 350) DEGs with a false discovery rate (FDR) p-value < 0.05 in SP+ vs. SP- cells depicted as red dots, including *SELE, GATA3, XCL2, IL7R, CD69, KLRD1, IL18R1, VCAM1, TNFRSF10A*, and *SPOCK2*. Grey dots represent genes with an FDR p-value > 0.05. (**E**) Boxplots of myometrium SP associated genes, *ABCB1* (log_2_FC= 1.9, FDR p= 2.8×10^-5^), *ABCC1* (log_2_FC= 0.7, FDR p= 5.9 x10^-1^), *ABCG2* (log_2_FC= 1.1, FDR p= 9.5×10^-2^), *ESR1* (log_2_FC= −1.1, FDR p= 2.3 x10^-1^) and *PGR* (log_2_FC= −1.5, FDR p= 6.4 x10^-2^) and (**F**) MSC associated genes, *SUSD2* (log_2_FC= −1, FDR p= 5.6×10^-1^), *MCAM* (log_2_FC= −1.5, FDR p= 3.9×10^-2^), *PDGFRβ* (log_2_FC= −0.9, FDR p= 4.9×10^-1^), *CSPG4* (log_2_FC= −1.5, FDR p= 3.9×10^-2^), *CD44* (log_2_FC= −0.5, FDR p= 8.3×10^-1^), *CD34* (log_2_FC= −0.2, FDR p= 4.9×10^-1^), and *ITGA6* (log_2_FC= 0.9, FDR p= 2.1×10^-1^) in the SP- (in grey) and SP+ (in red) cell population (n = 3), genes are expressed in log_2_CPM. ns = not significant, with FDR > 0.05.

### A putative MyoSPC cluster is determined by single cell RNA-seq

A total of 9,775 cells from 5 myometrium samples passed quality control with an average of 98.3% sequencing saturation, or approximately 512,000 reads per cell. Uniform Manifold Approximation and Projection (UMAP) of myometrial (n = 5) single cell RNA-seq (scRNA-seq) revealed 7 main cell clusters (**Fig 3A**) with similar cell distribution patterns across the five myometrial samples (**Fig S1A**). Cluster identities were assigned using the expression profiles of canonical markers for cell populations expected to be found in the myometrium (**Fig 3B**) (37,38), including 4 different smooth muscle cell types, vascular myocytes, myocytes, myofibroblasts, and fibroblast. The cell proportion of each identified clusters was similar across patients with these muscle cell types dominant **(Fig S1B)**. Four MSC markers, *SUSD2, MCAM, PDGFRβ* and *CSPG4*, were found highly expressed in the vascular myocyte cluster (**Fig 3C and 3D**), a common MSC niche (39,40). Immunofluorescence suggests that all 4 MSC markers were found surrounding the blood vessels in a separate set of myometrial samples (**Fig 3E**). Known MSC properties such as quiescence (G0) and the low gene regulation dynamics (41–43) were determined by the cell cycle score and the velocity of the scRNA-seq data, respectively. We identified a small group of cells within the vascular myocyte cluster in the G1/G0 phase (**Fig 4A**) using a computational assignment of cell-cycle stage (44). Cell velocity, which predicts the future state of individual cells using the RNA splicing information from each cell (45), showed that the same group of cells in G1/G0 phase in the vascular myocyte cluster are depicted with low velocity vectors (**Fig 4B**), indicating low levels of transcriptional changes, another characteristic of stem cells (46). We defined cells within the vascular myocyte cluster presenting with high expression of MSC markers in a G1/G0 phase and with low velocity as the “MyoSPC” cluster.

**Figure 3.**
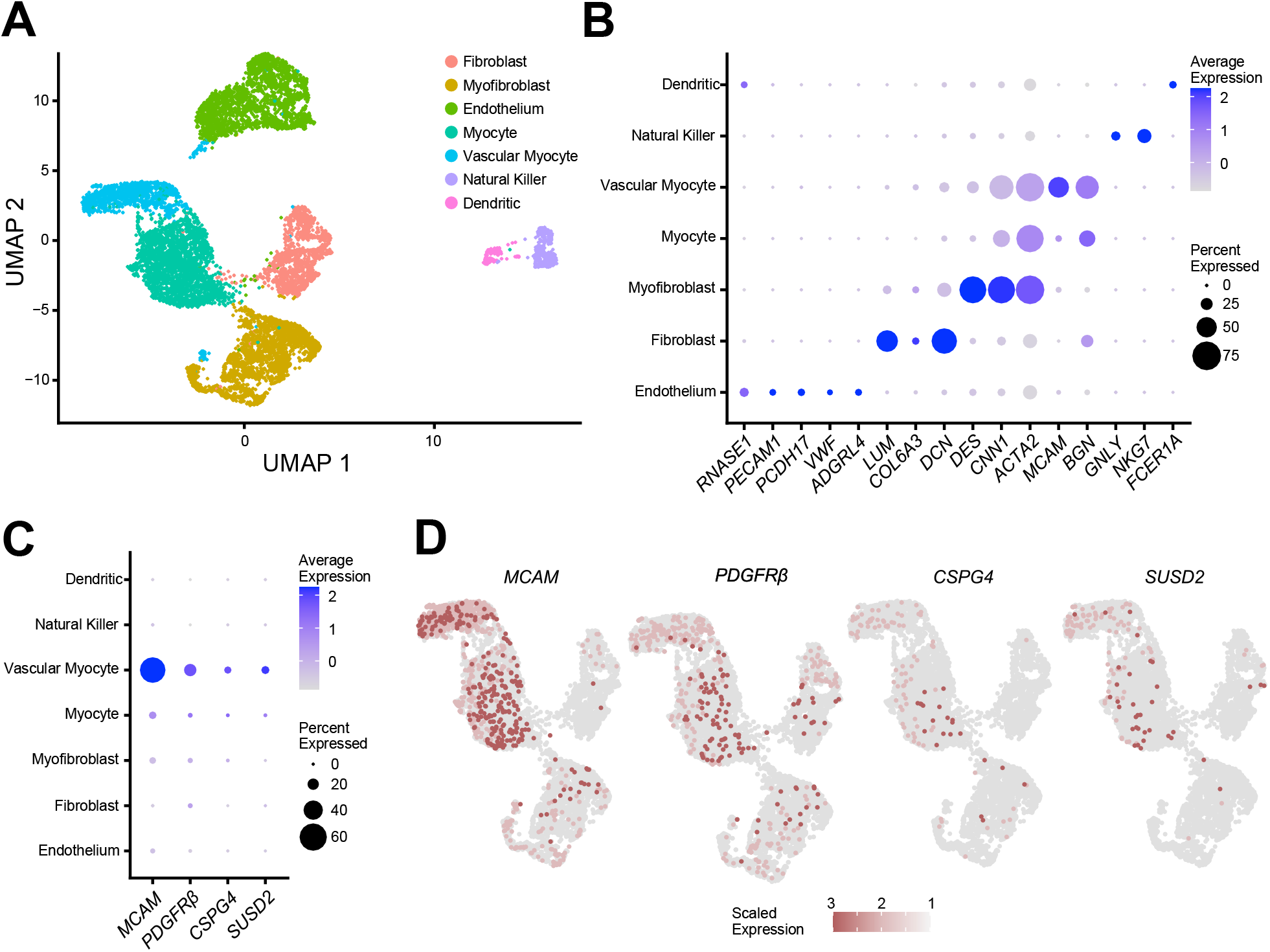
Single cell RNA-seq analysis of isolated cells from human myometrial samples. (**A**) Uniform manifold approximation and projection (UMAP) visualization of 9775 isolated cells from human myometrial samples (n = 5). Each cluster (n = 7) represent a cell population with a similar transcriptomic profile. (**B**) Dotplot for cluster identification using specific markers for cell types found in the myometrium. MSC marker gene expression in the different myometrial cell clusters shown in a dotplot (**C**) and by UMAP (**D**). Average gene expression and percentage of cells expressing the specific gene in each cell cluster are shown by the color intensity and the diameter of the dot, respectively, in B and C. Color gradient in the UMAP represents gene expression as log_2_CPM in D.

**Figure 4.**
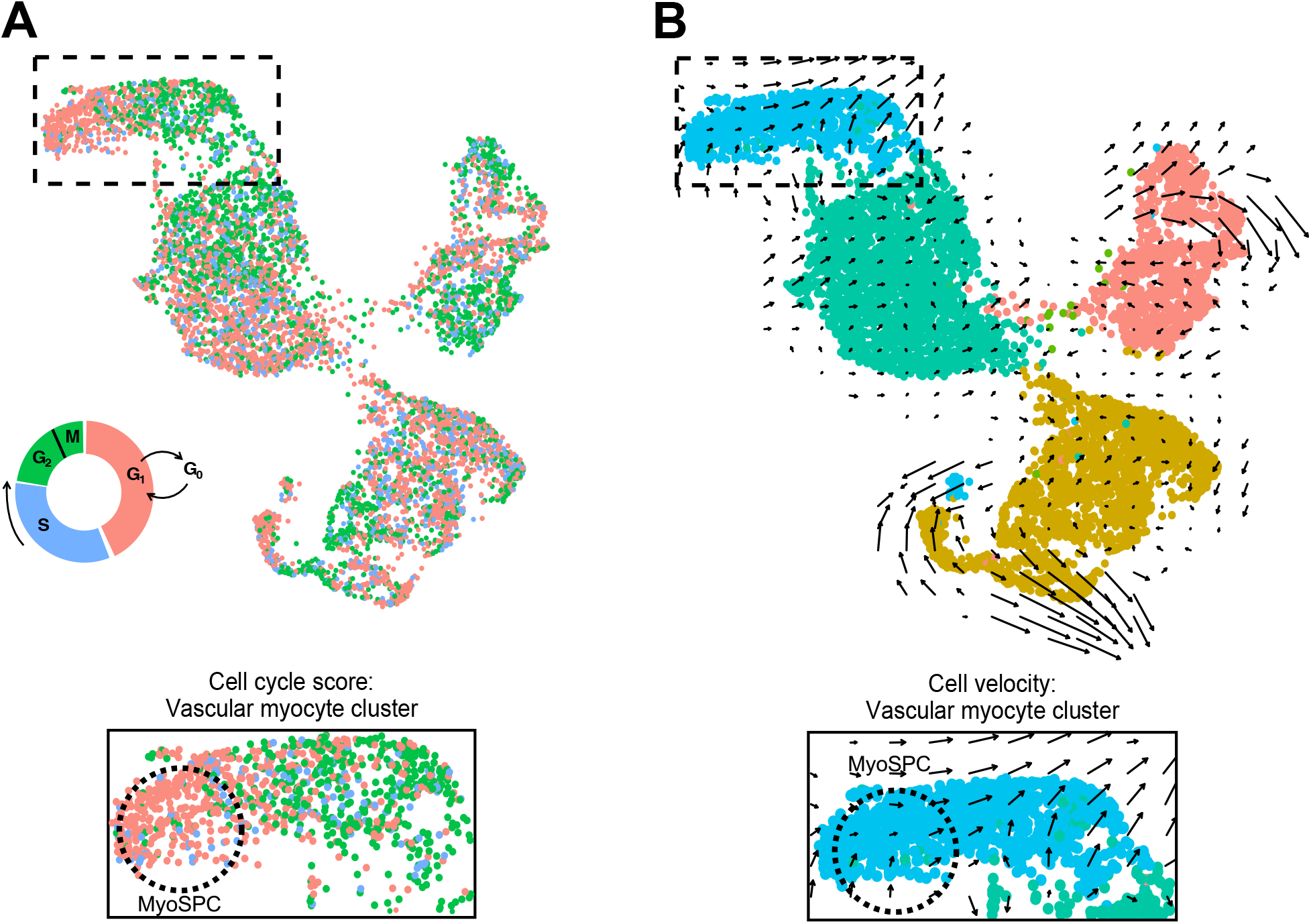
Identification of putative MyoSPCs from scRNA-seq. (**A**) Cell cycle score for myometrial cells visualized in the UMAP plot. Cells in G1/G0, S, and G2/M phases are plotted with the corresponding color. Boxed area is shown at higher magnification. (**B**) Cell velocity predicting the future state of individual cells illustrated in a UMAP plotted with the clusters as in panel A showed that in the vascular myocyte cluster, the same group of cells in G1/G0 phase exhibit low velocity. Boxed area is shown at higher magnification. Putative MyoSPCs are encircled with black dotted lines.

### Integrating bulk SUSD2+/^-^ RNA^-^seq and myometrial scRNA^-^seq reveals a new MyoSPC marker (CRIP1)

Transcriptomic analyses of SUSD2+/- bulk RNA and myometrial scRNA-seq were performed, and the results were integrated to discover possible overlapping MyoSPC markers. A total of 3,700 DEGs were found in the MyoSPC scRNA-seq cluster compared to the rest of the myometrial cells (**Table S3**). A little over half (1929 DEGs) of the MyoSPC DEGs overlapped significantly (p = 9.5 x 10^-81^) with the DEGs from the SUSD2 +/- bulk RNA-seq comparison (**Fig 5A and Table S4**). Correlation analysis of the log_2_ fold change (FC) in gene expression in the scRNA-seq analysis with the SUSD2+/- bulk RNA-seq confirmed that the MSC markers, *SUSD2, MCAM, PDGFRβ* and *CSPG4* were upregulated in both (**Fig 5B**). The most highly upregulated gene in the MyoSPC cluster, Cysteine-Rich Intestinal Protein 1 (*CRIP1*), is also significantly upregulated in the SUSD2+ cells (**Fig 5B**). UMAP plot showed that *CRIP1* was highly expressed in the vascular myocyte cluster (**Fig 5C**), and more particularly in the MyoSPC cluster (**Fig 5D**). *CRIP1* expression wasn’t differentially expressed (log_2_FC= − 0.2, FDR p= 9.9×10^-1^) in the RNA-seq results of the SP assay (**Fig S2A**). Although the cell distribution in each cluster was different **(Fig S2B and Table S5)**, we confirmed that *CRIP1* and the MSC markers were enriched in the MyoSPC cluster **(Fig S2C)** in cells from an orthogonal scRNA-seq study of myometrium from fibroid patients (38) when the cells were projected onto the UMAP shown in **Fig 3A**.

**Figure 5.**
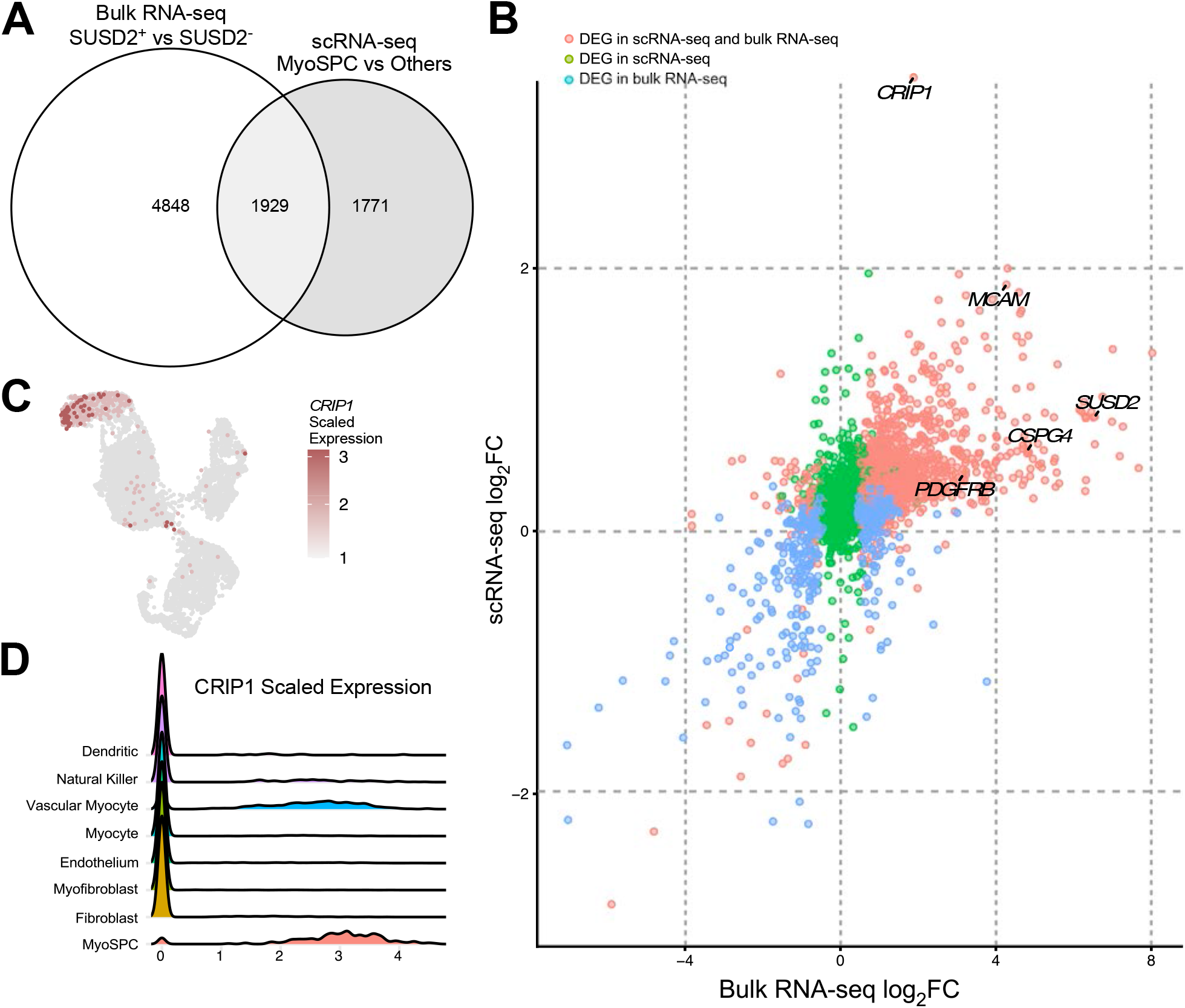
Integrated analysis of the bulk RNA-seq of SUSD2+ vs SUSD2- and the MyoSPC cluster vs the rest of the myometrial cells from the scRNA-seq. **(A)** Venn diagrams illustrate the overlapping DEGs between the bulk RNA-seq of SUSD2+ vs SUSD2- and the assigned MyoSPC cluster compared with the rest of the myometrial cells from the single cell RNA-seq analysis. **(B)** Scatter plot of Log_2_ fold change genes from bulk RNA-seq (x axis) and scRNA-seq (y axis). Non-significant genes in both analyses were represented in purple dots, and DEGs in the bulk RNA-seq only were represented in blue dots, DEGs in the scRNA-seq only were represented in green dots, and the DEGs in both analyses were represented in red dots. *CRIP1* is highly upregulated in the MyoSPC cluster (log_2_FC=3.1, adjusted p value = 4 x 10^-251^) and in the SUSD2+ cells (log_2_FC=1.9, FDR = 5 x 10^-4^). UMAP (**C**) and Ridge plots (**D**) display *CRIP1* scaled expression by cell cluster.

### CRIP1+ cells have common stem/progenitor cell properties

We next investigated the CRIP1 + cells to establish their stem cell bona fides. Immunofluorescence analysis using 3D imaging of the myometrial layer showed that CRIP1 + cells are located surrounding the PECAM1+ vascular endothelial cells, a common MSC niche (39,40,47) (**Fig 6A and Supplementary Video**). Interestingly, CRIP1 + cell immunofluorescence appeared to be predeominantly localized near the larger blood vessels and within a subset of SUSD2+ cells. Flow cytometry revealed that CRIP1 + cells represented between 2 to 5% of the total myometrial cells (**Fig 6B**). PECAM1 was used for negative selection of the smaller population of endothelial cells that also expressed CRIP1. CRIP1+/PECAM1- cells and the depleted cell population were sorted, and typical downstream stem cell assays were performed to determine if the CRIP1+/PECAM1- cells have stem/progenitor cell proprieties. Colony formation assays indicated that CRIP1+/PECAM1- sorted cells have a greater self-renewal capacity compared to the depleted sorted population (**Fig 6C**), with a significant increase of 4.5-fold greater number of colonies formed (**Fig 6D**), as well as a significant increase of the size of the colonies (**Fig 6E**). After 5 days in smooth muscle differentiation media, CRIP1+/PECAM1- cells were positive for ACTA2, indicating that they differentiated into smooth muscle cells (**Fig 6F**). Similarly, CRIP1+/PECAM1- cells were positive for Oil Red O staining (**Fig 6G**), and alkaline phosphatase activity (**Fig 6H**) when grown in either in adipogenic or osteogenic differentiation media, respectively, compared to CRIP1+/PECAM1- cells grown in control media, indicating that these putative MyoSPC cells have the capacity to differentiate into adipocytes and osteocytes.

**Figure 6.**
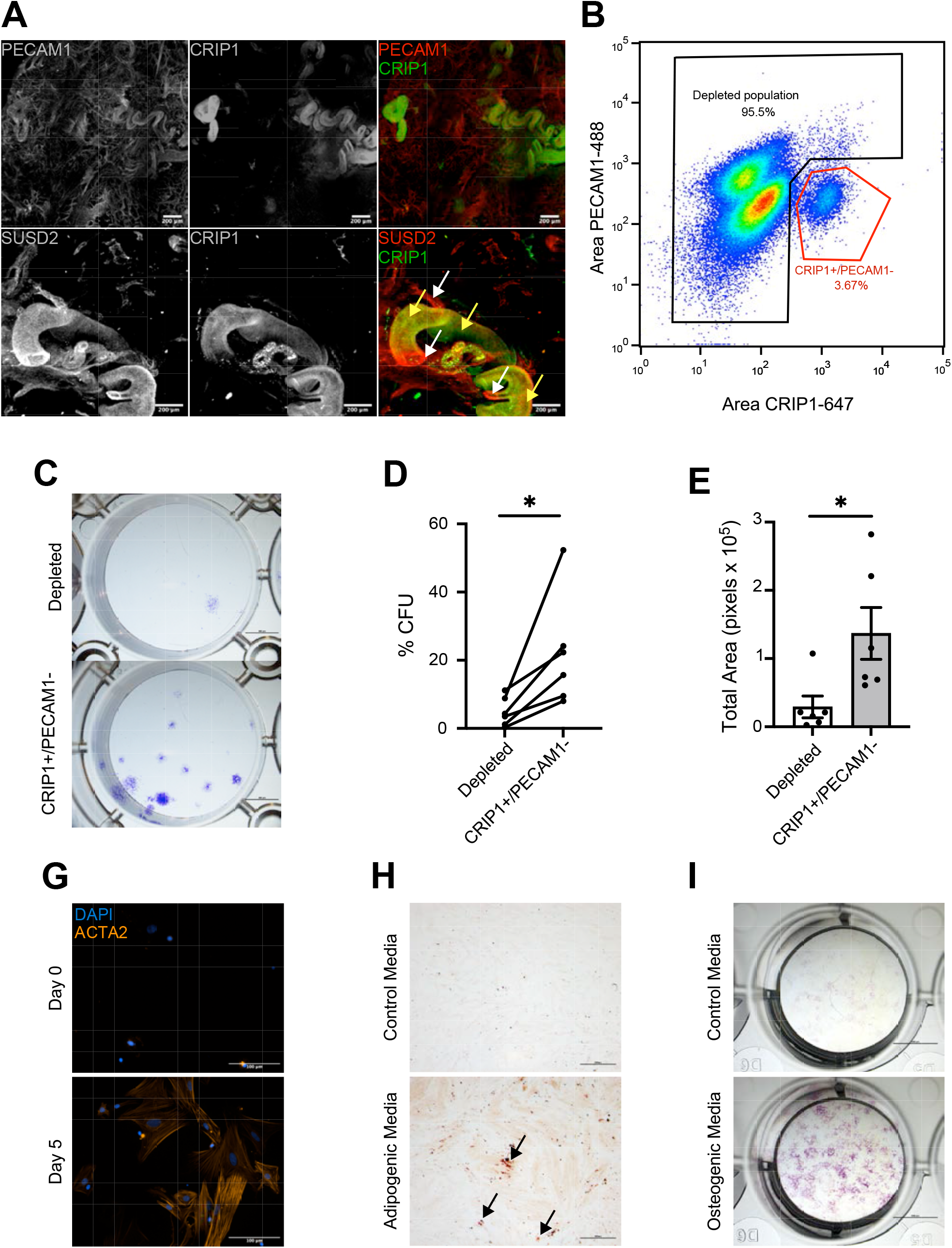
CRIP1^+^ cells have stem/progenitor cell characteristics. (**A**) Representative (n=3) immunofluorescence imaging of human myometrium using PECAM1 as an endothelial marker, SUSD2 as a mesenchymal stem cell marker, and CRIP1. Scale bar = 200 μm. (**B**) Representative (n=6) scatter plot of the gating strategy for CRIP1+/PECAM- cell sort. (**C**) Representative (n=6) images of colonies formed by the CRIP1+/PECAM- and depleted myometrial cells. (**D**) Plot of colony forming efficiency represented as %CFUs (#CFU/cells seeded × 100) of CRIP1+/PECAM- and depleted myometrial cells (n =6). (**E**) Total area of colony formed in pixels from CRIP1+/PECAM- and depleted myometrial cells (n =6). (**F**) ACTA2 immunofluorescence in CRIP1^+^/PECAM^-^ myometrial cells after differentiation. Scale bar = 100 μm. Representative (n=3) images of CRIP1+/PECAM- and depleted myometrial cells grown in control growth media and adipogenic **(G)** or osteogenic **(H)** differentiation media. Adipogenic and control cultures were stained with Oil Red O (red color, black arrows), and osteogenic and control cultures were stained for alkaline phosphatase activity (purple color). Scale bar for the adipogenic and osteogenic assays are 500 μm and 5 mm, respectively. *p<0.05; by student t test.

## Discussion

We have identified CRIP1 as a novel cell surface marker that enriches for a possible MyoSPC by combining the analyses of two next generation sequencing techniques, bulk RNA-seq from SUSD2+ and SUSD2- cells and scRNA-seq of total myometrial samples. Bulk RNA-seq, enriched for known MSC markers, but the large number of DEGs made it difficult to choose putative novel myometrium stem cell markers for further study. To reduce the number of candidate markers for validation and follow up studies, we used scRNA-seq to identify possible stem cells based on MSC markers. Subsequent stem cell assays confirmed that CRIP1 + cells have MSC properties, and further studies are underway to determine whether these cells could be a cell of origin for uterine fibroids.

CRIP1, Cysteine-rich intestinal protein 1, is a member of the LIM/double zincfinger proteins that is predicted to be a novel biomarker in multiple cancers and can promote several biological processes, including cell migration, invasion and epithelial-mesenchymal transition by activating Wnt/β□catenin signaling, an essential pathway that maintains stem cell homeostasis in many tissues (48–50). In an earlier study of SP+/- cells in fibroids (51), *CRIP1* was among the DEGs detected by microarray analysis. In that study, the authors demonstrated that the SP+ cells from fibroid tissues have stem cell characteristics, including self-renewal and differentiation into adipose and osteocyte cells. In the present study, we used myometrial samples from non-fibroid patients (MyoN) samples because we recently reported that myometria from fibroid patients (MyoF) have a different transcriptomic profile compared to MyoN samples, including an enrichment of DEGs in a leiomyoma disease ontology panel (52). Here we have reported that *CRIP1* expression was not differentially expressed and that the expression of the MSC markers, *SUSD2, MCAM, PDGFRβ*, and *CSPG4* were decreased in SP+ compared to the SP- from MyoN cells. These discordant results could arise from the tissue type, that is, fibroid tumor versus MyoN, or because the SP technique could be more applicable to other tissues (20) or hematopoietic stem cells (HSCs). It is worth noting that HSC exhibit a specific ABC transporter gene expression profile distinct from other stem cells, including MSCs (53). HSC expressed higher level of most of the ABC transporters including *ABCB1, ABCC1* and *ABCG2* compared to other stem cells. Indeed, the SP+ MyoN cells were enriched for ABC transporters and HSC-associated genes. Additionally, the SP technique relies on an intact cell metabolism and considerable variation in results has been observed (20).

Our scRNA-seq results suggested that human myometrium has at least 7 different cell types, including different types of smooth muscle cells, endothelial cells, and immune cells. Similar clusters were reported in a scRNA-seq comparison of fibroids and myometrium (38). The depth of sequencing for each cell was close to saturation allowing us to identify a small cell population with stem cell characteristics, including the expression of MSC markers *SUSD2, MCAM, PDGFRβ*, and *CSPG4*, a quiescent (G0) cell cycle state (41) and low transcriptomic activity/low RNA velocity (41–43). CRIP1+/PECAM1- cells were primarily located in the perivascular region, a common MSC niche (39,40,47), particularly by the larger myometrial blood vessels. These results were consistent with our scRNA-seq results showing that *CRIP1* expression was most highly expressed in the vascular myocyte cluster. Moreover, immunofluorescence staining showed that CRIP1 + cells were a subset the SUSD2+ cells and that CRIP1+/PECAM1- cells account for only 2 to 5% of the total human myometrial cells, a typical stem cell proportion in adult tissues (54). Interestingly, the depleted population was able to form a few colonies, an indication that some cells in the depleted population also have self-renewal properties. Similar results were observed by us using other MSC markers to isolate myometrium stem/progenitor cells, including SUSD2, MCAM, or PDGFRβ (18). This finding suggests that further enrichment of the MyoSPCs with some of the other MSC markers is possible or that myometrial cell plasticity is more common than heretofore appreciated.

In summary, we have identified CRIP1 as a novel marker of MyoSPCs from integration of two transcriptome sequencing techniques, sorted bulk cell and single cell RNA-seq. Induction of a known fibroid subtype mutation in CRIP1+/PECAM1- cells and their subsequent development into fibroid-like cells, could advance our understanding of fibroid etiology based on the hypothesis that a dysregulated MyoSPC is the origin of uterine fibroids.

### Study approval

The use of human tissue specimens was approved by the Spectrum Health Systems and Michigan State University Institutional Review Boards (MSU IRB Study ID: STUDY00003101, SR IRB #2017-198) as secondary use of biobank materials.

## Author contributions

Experimental design (E.N.P, J.M.T) collected data and performed experiments (E.N.P, T.J.C, S.F), analyzed data (E.N.P, R.S, K.H.L, R.A, J.M.T), wrote/reviewed manuscript (E.N.P, T.J.C, S.F, R.S, K.H.L, R.A, J.M.T).

## Acknowledgments

We would like to thank the patients who consented for the study, the Spectrum Health Systems Universal Biorepository staff, and the Van Andel Institute Genomics Core (RRID:SCR_022913), especially Marie Adams, Rebecca Siwicki and Julie Koeman, for their assistance with the construction and sequencing of 10X libraries for the single cell RNA-seq and bulk RNA-seq. This work was supported by grants HD096259 and HD100959 from the Eunice Kennedy Shriver National Institute of Child Health and Human Development (to J.M.T) and the SRI/Bayer Discovery/Innovation Grant (to E.N.P).

## Methods

### Sample collection and cell isolation

The use of human tissue specimens was approved by the Spectrum Health Systems Institutional Review Board as secondary use of biobank materials. Myometrial samples from non-fibroid patients (MyoN) were obtained following total hysterectomy from premenopausal (aged 34-50), self-identified Caucasian women. No fibroids were detected by ultrasound prior to surgery. All patients who participated in the study gave consent to donate tissue through the Spectrum Health Biorepository. Myometrial samples were washed with PBS, dissected away from non-myometrial tissue, and minced. Cells were isolated by incubation at 37°C in baffled flasks containing digestion media (DMEM/F12, 10% fetal bovine serum (FBS), collagenase type I, DNAse type I, and MgCl_2_) with agitation. The resulting cell suspensions were strained through 100- and 40-μm cell strainers, washed with warm media (DMEM/F12 containing 15% FBS),and centrifuged. Isolated cells were then stored in freeze media (90% FBS, 10% DMSO) at −80 °C until needed.

### Cell staining for FACS

Human primary myometrial cells were thawed and resuspended in 1% bovine serum albumin blocking buffer for 20 min at room temperature (RT). Cells were then incubated with the primary antibody for 45 min at RT; SUSD2-PE anti-human (Miltenyi Biotec, #130-117-682), PECAM1-FITC anti-human (ThermoFisher, #11-0319-42), and CRIP1 rabbit anti-human (ThermoFisher, #PA5-24643). For CRIP1/PECAM1 staining, cells were incubated with an Alexa-647 anti-rabbit secondary antibody for 30 min at RT. Stained myometrial cells were then wash with flow buffer and resuspend in 1 mL of flow buffer with 1 μg of 4’,6-diamidino-2-phenylindole (DAPI) or Propidium Iodode (PI), depending on the experiment, for live dead discrimination. Cells were sorted by the flow cytometry core at Van Andel Research Institute (VARI) using a FACSymphony S6 cytometer (BD Biosciences) and analyzed with FlowJo Software (BD Biosciences, version 10.8.1).

The side population assay was conducted as described previously (17). Briefly, live cells were incubated with 5 μg/mL of Hoechst 33342 dye for 90 min. As a negative control, separate aliquots of cells from the same patients were treated with 25 μg/ml of verapamil (Sigma) prior to addition of the Hoechst dye. PI was added to stained cells with and without verapamil treatment and analyzed in a MoFlo Astrios (Beckman Coulter) for side population gating by Hoechst red and blue filters and sorting in media (DMEM/F12) at 4 °C.

### RNA Isolation, Library Preparation and Sequencing

Total RNA was isolated from sorted cells using an RNeasy mini kit (Qiagen) and stored at −80 °C in nuclease-free water. RNA integrity values were determined with an Agilent 2100 Bioanalyzer (ThermoFisher), and values ≥7.5 were used for library preparation and paired-end (2 × 100 bp) RNA-sequencing on an Illumina NextSeq 6000 instrument (Illumina). Libraries were prepared using a Kapa RNA HyperPrep kit with ribosomal reduction, pooled, and sequenced on flowcells to yield approximately 50–60 million reads/sample. Raw fastq files were deposited in the NCBI Gene Expression Omnibus (GSEXXXX).

For single cell RNA-seq, dead cells were removed from digested myometrial cells using the Dead Cell Removal Kit (Miltenyi Biotec, #130-090-101) per manufacturer’s instructions. Live myometrial cells from 5 non-fibroid patients were then sequenced. Libraries were generated and sequenced using the 10X Chromium Next GEM Single Cell 3□ GEM kit (10X Genomics, v2) platform according to the manufacturer’s instructions. 2 x 75 bp, paired end sequencing was performed on an Illumina NovaSeq 6000 sequencer using an S2 flow cell, 100 cycle sequencing kit (v1.5) to a minimum depth of 50K reads per cell (Illumina Inc., San Diego, CA, USA). Base calling was done by Illumina RTA3 and output was demultiplexed and converted to FastQ format with Cell Ranger (10X Genomics, v3.1.0). Raw fastq files were deposited in the NCBI Gene Expression Omnibus (GSEXXXX).

### RNA Seq Analysis

For bulk RNA-seq, reads were trimmed for quality and adapters using TrimGalore (version 0.6.5), and quality trimmed reads were assessed with FastQC (version 0.11.7). Trimmed reads were mapped to Homo sapiens genome assembly GRCh38 (hg38) using STAR (version 2.7.9a). Reads overlapping Ensembl annotations (version 99) were quantified with STAR prior to model-based differential expression analysis using the edgeR-robust method with paired samples. Genes with low counts per million (CPM) were removed using the filterByExpr function from edgeR (55). Scatterplots of two selected principal components was constructed with the PCAtools R package (version 2.5.13) to verify sample separation prior to statistical testing. Generalized linear models were used to determine if principal components were significantly associated with cell type. Genes were considered differentially expressed if their respective edgeR- robust FDR corrected p-values were less than 0.05. Differential expression was calculated by comparing SUSD2+ to SUSD2- cells. DEGs were visualized with volcano plots and heatmaps generated using the EnhancedVolcano (version 1.8.0) and pheatmap (version 1.0.12) packages in R. Box plots of the log_2_(CPM) values were generated using the R package ggplot2 (version 3.4.0).

For scRNA-seq, demultiplexed sequencing reads were processed and aligned to the *Homo sapiens* genome assembly GRCh38 (hg38) using STAR (version 2.7.9a) with 10X Genomics Cell Ranger (version 3.1.0). Samples were merged using the integration anchors function of the Seurat package (version 4.2.1) from R (56). Genes expressed in fewer than three cells in a sample were excluded, as well as cells that expressed fewer than 200 genes and mitochondrial gene content >5% of the total unique molecular identifier count. Data were normalized using a global-scaling normalization method (56) that normalizes the feature expression measurements for each cell by the total expression, multiplies this by a scale factor (10,000), and then log-transforms the results. The top 2,000 most variable genes that were used for cell clustering were found using the *FindVariableFeatures* function and were then normalized using the *ScaleData* function. Based on an elbow plot generated using the *Elbowplot* function of Seurat, we selected 15 principal components (PC) for downstream analyses. Cell clusters were generated using *FindNeighbors* and *FindClusters* functions. For visualization, UMAPs were generated using the *RunUMAP, FeaturePlot* and *DimPlot* functions. The *DotPlot* Seurat function was used to generate dot plots to visualize gene expression for each assigned cluster. Cell cycle score and velocity were determined using the functions *CellCycleScoring* from Seurat and *RunVelocity* from SeuratWrappers (version 0.3.0), respectively. The “stem cell” cluster was selected using the *CellSelector* function from Seurat. The *RidgePlot* function from the Seurat R package was used for the visualization of *CRIP1* gene expression in the different myometrium clusters.

Venn diagrams of the overlapping DEGs from the bulk RNA-seq of SUSD2+ and SUSD2- cells and DEGs of scRNA-seq of the “MyoSPC” cluster compared to the rest of the myometrial cells were constructed using the eulerr package (version 6.1.1). The scatter plot of overlapping DEG from bulk- and scRNA-seq was generated with ggplot2 (version 3.4.0). *CRIP1* expression in the MyoSPC cluster was confirmed using a single cell data from the NCBI Gene Expression Omnibus (GSE162122). Single cell data from 5 MyoF from that study were mapped to the hg38 using STAR. A total of 18939 cells passed quality control and were projected onto our reference UMAP using the function MapQuery from Seurat.

### Imaging

Whole mount immunofluorescent staining (57) was performed for human myometrial samples that were fixed in a 4:1 solution of methanol:DMSO. Tissue was removed from the fixative, rehydrated in (1:1) methanol: PBST (PBS, 1% triton) solution and washed in 100% PBST. Samples were incubated in a blocking solution (PBS, 2% powdered milk, 1% triton) and then stained with a 1:500 dilution for primary antibodies in blocking solution for 7 nights at 4°C. Primary antibodies used were Rabbit anti-human CRIP1 (ThermoFisher, #PA5-24643), Mouse anti-human SUSD2 (Biolegend #327401) and Mouse anti-human PECAM1 (Abcam, #ab9498). Samples were then incubated with secondary antibodies, Donkey anti-Rabbit IgG Alexa Fluor 555 (Invitrogen, #A31572), Goat anti-Mouse IgG Alexa Fluor 647 (Invitrogen, #A21235) at a dilution of 1:300 and Hoechst dye, for three nights at 4°C. Samples were transferred to a methanol: PBST (1:1) solution, then washed in methanol and incubated at 4°C overnight in a 3% H_2_O_2_ solution diluted in methanol. Tissues were then washed in methanol and cleared in benzyl alcohol: benzyl benzoate (1:2) overnight. Imaging was performed using a Leica SP8 TCS white light laser confocal microscope utilizing 10x air objective and a 7.0 μm Z stack (58). Imaris v9.2.1 (Bitplane) commercial software was used to analyze confocal image files and create 3D renderings. The image files were imported into Imaris 3D surpass mode and 3D renderings were created using the Surface plugin. Images were captured using the Snapshot plugin of Imaris and video were generated using the Animation plugin.

### Colony formation and mesenchymal lineage differentiation

CRIP1+/PECAM1- and the depleted cell populations were sorted as described above and plated at 50 cells/cm^2^ in triplicate in growth media (DMEM/F12, 10% FBS) overnight, then grown in MesenPro RS (Thermo Fisher, # 12746012) for 2 to 3 weeks. Cultures were fixed in 4% paraformaldehyde (PFA) and stained with crystal violet for colony visualization. Colonies were counted and total surface area was estimated using ImageJ (version 1.53k), and the percent colony-forming units (CFUs) was calculated as (number of colonies/number of cells plated) × 100 and averaged for triplicates. Matched CRIP1+/PECAM1- and depleted populations were cultured in different wells of the same plate, and both cell types were assayed on the same day. Images were taken using a Nikon SMZ18 microscope and Ds-Ri1 camera (Nikon Instruments Inc.). For osteogenic and adipogenic differentiation, CRIP1+/PECAM1- cells were plated in triplicate at 50% to 80% confluency in growth media (DMEM/F12, 10% FBS) overnight and then cultured for 10 days in fresh StemPro Adipogenesis Differentiation (Thermo Fisher Scientific, #A1007001) or StemPro Osteogenesis Differentiation (Thermo Fisher Scientific, #A1007201) media according to the manufacturer’s instructions. Cells were cultured in regular growth media to serve as differentiation controls. To assay adipogenic differentiation, cultures were fixed in 4% PFA and stained using Oil Red O (Sigma, #01391) according to the manufacturer’s instructions. To assay osteogenic differentiation, cultures were stained for alkaline phosphatase activity using the Alkaline Phosphatase (AP), Leukocyte kit (Sigma, #86R-1KT) according to the manufacturer’s instructions. For smooth muscle differentiation, cells were plated on 1 mg/ml dried rat tail collagen (Corning, #354 236) in 8 well chamber slides with growth media (DMEM/F12, 10% FBS) overnight and then cultured in Medium 231 with a smooth muscle differentiation supplement (Thermo Fisher, #M231500). Cells were fixed in 4% PFA at the indicated times (D0: before adding the differentiation media, D5: 5 days in differentiation media) and stained using αSMA-Cy3 (Sigma, #C6198). Images were taken using a Nikon Eclipse Ni-U or Nikon SMZ18 microscope and Ds-Qi1MC or Ds-Ri1 camera (Nikon Instruments Inc.).

### Statistical analyses

Bioinformatic statistics were performed using the listed packages in R (version 4.0.2). DEGs of the bulk RNA-seq from SUSD2+ vs. SUSD2- and the SP+ vs. SP- were identified as those having an Benjamini–Hochberg FDR corrected p <0.05 (59). Data with unequal variances were log transformed, and homogeneity of variances verified before completion of analyses. DEGs of the scRNA-seq were calculated using the non-parametric Wilcoxon Rank Sum test. Adjusted p-value, based on Bonferroni correction using all features in the dataset was used to determine significance. DEGs with adjusted p-value < 0.05 were consider as significant. Hypergeometric testing was performed using the function phyper from R. For the colony formation assays, comparison of two means was performed with a two-sided student t test, and significance was determined at p <0.05 after confirming normal distribution using Graphpad Prism (version 9.4.1).

---

**Figure S1. Cell distribution across cell clusters in the single cell RNA-seq.** (**A**) Uniform manifold approximation and projection (UMAP) visualization of 9775 isolated cells from human myometrial samples (n = 5). Each color dot represents cells from a myometrials from a different patient. UMAP plot shows that each patient’s cells are well distributed across clusters. (**B**) Cell proportion of each cluster as a percentage across patients.

**Figure S2. *CRIP1* expression in the side population and an orthogonal single cell study.** (**A**) *CRIP1* expression in log_2_CPM of the RNA-seq results from the SP+ is not significantly different from that of the SP- cells (FDR>0.05). (**B**) Projection of a data set of 18,939 cells from 5 myometrial samples from fibroids patients (38) onto the UMAP in Fig 3A. (**C**) Dotplot of mesenchymal stem cell markers and *CRIP1* gene expression in the different myometrial cell clusters as defined in Fig 3C.

## Notes

### Competing Interest Statement

The authors have declared no competing interest.

### Summary of Updates

This version has been updated with new findings and updated results

## References

1. Zimmermann A, Bernuit D, Gerlinger C, Schaefers M, Geppert K. Prevalence, symptoms and management of uterine fibroids: an international internet-based survey of 21,746 women. BMC Womens Health. 2012;12:6.

2. Gupta S, Jose J, Manyonda I. Clinical presentation of fibroids. Best Pract Res Clin Obstet Gynaecol. 2008;22(4):615–626.

3. Farris M, Bastianelli C, Rosato E, Brosens I, Benagiano G. Uterine fibroids: an update on current and emerging medical treatment options. Ther Clin Risk Manag. 2019;15:157–178.

4. Sohn GS, Cho S, Kim YM, Cho CH, Kim MR, Lee SR, Working Group of Society of Uterine L. Current medical treatment of uterine fibroids. Obstet Gynecol Sci. 2018;61(2):192–201.

5. Ono M, Maruyama T. Stem Cells in Myometrial Physiology. Semin Reprod Med. 2015;33(5):350–356.

6. Teixeira J, Rueda BR, Pru JK. Uterine stem cells. StemBook. Cambridge (MA) 2008.

7. Canevari RA, Pontes A, Rosa FE, Rainho CA, Rogatto SR. Independent clonal origin of multiple uterine leiomyomas that was determined by X chromosome inactivation and microsatellite analysis. Am J Obstet Gynecol. 2005;193(4):1395–1403.

8. Zhang P, Zhang C, Hao J, Sung CJ, Quddus MR, Steinhoff MM, Lawrence WD. Use of X-chromosome inactivation pattern to determine the clonal origins of uterine leiomyoma and leiomyosarcoma. Hum Pathol. 2006;37(10):1350–1356.

9. Holdsworth-Carson SJ, Zaitseva M, Vollenhoven BJ, Rogers PA. Clonality of smooth muscle and fibroblast cell populations isolated from human fibroid and myometrial tissues. Mol Hum Reprod. 2014;20(3):250–259.

10. Fialkow PJ. Clonal origin of human tumors. Annu Rev Med. 1979;30:135–143.

11. Mas A, Nair S, Laknaur A, Simon C, Diamond MP, Al-Hendy A. Stro-1/CD44 as putative human myometrial and fibroid stem cell markers. Fertil Steril. 2015;104(1):225–234 e223.

12. Yin P, Ono M, Moravek MB, Coon JSt, Navarro A, Monsivais D, Dyson MT, Druschitz SA, Malpani SS, Serna VA, Qiang W, Chakravarti D, Kim JJ, Bulun SE. Human uterine leiomyoma stem/progenitor cells expressing CD34 and CD49b initiate tumors in vivo. J Clin Endocrinol Metab. 2015;100(4):E601–606.

13. Bulun SE. Uterine fibroids. N Engl J Med. 2013;369(14):1344–1355.

14. Szotek PP, Chang HL, Zhang L, Preffer F, Dombkowski D, Donahoe PK, Teixeira J. Adult mouse myometrial label-retaining cells divide in response to gonadotropin stimulation. Stem Cells. 2007;25(5):1317–1325.

15. Patterson AL, George JW, Chatterjee A, Carpenter T, Wolfrum E, Pru JK, Teixeira JM. Label-Retaining, Putative Mesenchymal Stem Cells Contribute to Myometrial Repair During Uterine Involution. Stem cells and development. 2018.

16. Ono M, Maruyama T, Masuda H, Kajitani T, Nagashima T, Arase T, Ito M, Ohta K, Uchida H, Asada H, Yoshimura Y, Okano H, Matsuzaki Y. Side population in human uterine myometrium displays phenotypic and functional characteristics of myometrial stem cells. Proc Natl Acad Sci U S A. 2007;104(47):18700–18705.

17. Chang HL, Senaratne TN, Zhang L, Szotek PP, Stewart E, Dombkowski D, Preffer F, Donahoe PK, Teixeira J. Uterine leiomyomas exhibit fewer stem/progenitor cell characteristics when compared with corresponding normal myometrium. Reprod Sci. 2010;17(2):158–167.

18. Patterson AL, George JW, Chatterjee A, Carpenter TJ, Wolfrum E, Chesla DW, Teixeira JM. Putative human myometrial and fibroid stem-like cells have mesenchymal stem cell and endometrial stromal cell properties. Human reproduction. 2020;35(1):44–57.

19. Ono M, Kajitani T, Uchida H, Arase T, Oda H, Uchida S, Ota K, Nagashima T, Masuda H, Miyazaki K, Asada H, Hida N, Mabuchi Y, Morikawa S, Ito M, Bulun SE, Okano H, Matsuzaki Y, Yoshimura Y, Maruyama T. CD34 and CD49f Double-Positive and Lineage Marker-Negative Cells Isolated from Human Myometrium Exhibit Stem Cell-Like Properties Involved in Pregnancy-Induced Uterine Remodeling. Biol Reprod. 2015;93(2):37.

20. Golebiewska A, Brons NH, Bjerkvig R, Niclou SP. Critical appraisal of the side population assay in stem cell and cancer stem cell research. Cell Stem Cell. 2011;8(2):136–147.

21. Masuda H, Anwar SS, Buhring HJ, Rao JR, Gargett CE. A novel marker of human endometrial mesenchymal stem-like cells. Cell Transplant. 2012;21(10):2201–2214.

22. Redvers RP, Li A, Kaur P. Side population in adult murine epidermis exhibits phenotypic and functional characteristics of keratinocyte stem cells. Proc Natl Acad Sci USA. 2006;103(35):13168–13173.

23. Schienda J, Engleka KA, Jun S, Hansen MS, Epstein JA, Tabin CJ, Kunkel LM, Kardon G. Somitic origin of limb muscle satellite and side population cells. Proc Natl Acad Sci USA. 2006;103(4):945–950.

24. Fox JC, Nakayama T, Tyler RC, Sander TL, Yoshie O, Volkman BF. Structural and agonist properties of XCL2, the other member of the C-chemokine subfamily. Cytokine. 2015;71(2):302–311.

25. Lopez-Cabrera M, Santis AG, Fernandez-Ruiz E, Blacher R, Esch F, Sanchez-Mateos P, Sanchez-Madrid F. Molecular cloning, expression, and chromosomal localization of the human earliest lymphocyte activation antigen AIM/CD69, a new member of the C-type animal lectin superfamily of signal-transmitting receptors. J Exp Med. 1993;178(2):537–547.

26. Akashi K, Kondo M, von Freeden-Jeffry U, Murray R, Weissman IL. Bcl-2 rescues T lymphopoiesis in interleukin-7 receptor-deficient mice. Cell. 1997;89(7):1033–1041.

27. Bongen E, Vallania F, Utz PJ, Khatri P. KLRD1-expressing natural killer cells predict influenza susceptibility. Genome Med. 2018;10(1):45.

28. Garlanda C, Dinarello CA, Mantovani A. The interleukin-1 family: back to the future. Immunity. 2013;39(6):1003–1018.

29. Passarelli C, Selvatici R, Carrieri A, Di Raimo FR, Falzarano MS, Fortunato F, Rossi R, Straub V, Bushby K, Reza M, Zharaieva I, D’Amico A, Bertini E, Merlini L, Sabatelli P, Borgiani P, Novelli G, Messina S, Pane M, Mercuri E, Claustres M, Tuffery-Giraud S, Aartsma-Rus A, Spitali P, T’Hoen PAC, Lochmuller H, Strandberg K, Al-Khalili C, Kotelnikova E, Lebowitz M, Schwartz E, Muntoni F, Scapoli C, Ferlini A. Tumor Necrosis Factor Receptor SF10A (TNFRSF10A) SNPs Correlate With Corticosteroid Response in Duchenne Muscular Dystrophy. Front Genet. 2020;11:605.

30. Liu G, Ren F, Song Y. Upregulation of SPOCK2 inhibits the invasion and migration of prostate cancer cells by regulating the MT1-MMP/MMP2 pathway. PeerJ. 2019;7:e7163.

31. Nabors LK, Wang LD, Wagers AJ, Kansas GS. Overlapping roles for endothelial selectins in murine hematopoietic stem/progenitor cell homing to bone marrow. Exp Hematol. 2013;41(7):588–596.

32. Tufa DM, Yingst AM, Shank T, Shim S, Trahan GD, Lake J, Woods R, Jones KL, Verneris MR. Transient Expression of GATA3 in Hematopoietic Stem Cells Facilitates Helper Innate Lymphoid Cell Differentiation. Front Immunol. 2019;10:510.

33. Bujanover N, Goldstein O, Greenshpan Y, Turgeman H, Klainberger A, Scharff Y, Gazit R. Identification of immune-activated hematopoietic stem cells. Leukemia. 2018;32(9):2016–2020.

34. Pinho S, Wei Q, Maryanovich M, Zhang D, Balandran JC, Pierce H, Nakahara F, Di Staulo A, Bartholdy BA, Xu J, Borger DK, Verma A, Frenette PS. VCAM1 confers innate immune tolerance on haematopoietic and leukaemic stem cells. Nat Cell Biol. 2022;24(3):290–298.

35. Maleki M, Ghanbarvand F, Reza Behvarz M, Ejtemaei M, Ghadirkhomi E. Comparison of mesenchymal stem cell markers in multiple human adult stem cells. Int J Stem Cells. 2014;7(2):118–126.

36. Patterson AL, George JW, Chatterjee A, Carpenter T, Wolfrum E, Pru JK, Teixeira JM. Label-Retaining, Putative Mesenchymal Stem Cells Contribute to Murine Myometrial Repair During Uterine Involution. Stem Cells Dev. 2018;27(24):1715–1728.

37. Pique-Regi R, Romero R, Garcia-Flores V, Peyvandipour A, Tarca AL, Pusod E, Galaz J, Miller D, Bhatti G, Para R, Kanninen T, Hadaya O, Paredes C, Motomura K, Johnson JR, Jung E, Hsu CD, Berry SM, Gomez-Lopez N. A single-cell atlas of the myometrium in human parturition. JCI Insight. 2022;7(5).

38. Goad J, Rudolph J, Zandigohar M, Tae M, Dai Y, Wei JJ, Bulun SE, Chakravarti D, Rajkovic A. Single-cell sequencing reveals novel cellular heterogeneity in uterine leiomyomas. Hum Reprod. 2022;37(10):2334–2349.

39. Crisan M, Yap S, Casteilla L, Chen CW, Corselli M, Park TS, Andriolo G, Sun B, Zheng B, Zhang L, Norotte C, Teng PN, Traas J, Schugar R, Deasy BM, Badylak S, Buhring HJ, Giacobino JP, Lazzari L, Huard J, Peault B. A perivascular origin for mesenchymal stem cells in multiple human organs. Cell Stem Cell. 2008;3(3):301–313.

40. Bautch VL. Stem cells and the vasculature. Nat Med. 2011;17(11):1437–1443.

41. O’Connor SA, Feldman HM, Arora S, Hoellerbauer P, Toledo CM, Corrin P, Carter L, Kufeld M, Bolouri H, Basom R, Delrow J, McFaline-Figueroa JL, Trapnell C, Pollard SM, Patel A, Paddison PJ, Plaisier CL. Neural G0: a quiescent-like state found in neuroepithelial-derived cells and glioma. Mol Syst Biol. 2021;17(6):e9522.

42. Riba A, Oravecz A, Durik M, Jimenez S, Alunni V, Cerciat M, Jung M, Keime C, Keyes WM, Molina N. Cell cycle gene regulation dynamics revealed by RNA velocity and deeplearning. Nat Commun. 2022;13(1):2865.

43. Zywitza V, Misios A, Bunatyan L, Willnow TE, Rajewsky N. Single-Cell Transcriptomics Characterizes Cell Types in the Subventricular Zone and Uncovers Molecular Defects Impairing Adult Neurogenesis. Cell Rep. 2018;25(9):2457–2469 e2458.

44. Scialdone A, Natarajan KN, Saraiva LR, Proserpio V, Teichmann SA, Stegle O, Marioni JC, Buettner F. Computational assignment of cell-cycle stage from single-cell transcriptome data. Methods. 2015;85:54–61.

45. La Manno G, Soldatov R, Zeisel A, Braun E, Hochgerner H, Petukhov V, Lidschreiber K, Kastriti ME, Lonnerberg P, Furlan A, Fan J, Borm LE, Liu Z, van Bruggen D, Guo J, He X, Barker R, Sundstrom E, Castelo-Branco G, Cramer P, Adameyko I, Linnarsson S, Kharchenko PV. RNA velocity of single cells. Nature. 2018;560(7719):494–498.

46. Efroni S, Duttagupta R, Cheng J, Dehghani H, Hoeppner DJ, Dash C, Bazett-Jones DP, Le Grice S, McKay RD, Buetow KH, Gingeras TR, Misteli T, Meshorer E. Global transcription in pluripotent embryonic stem cells. Cell Stem Cell. 2008;2(5):437–447.

47. Lv FJ, Tuan RS, Cheung KM, Leung VY. Concise review: the surface markers and identity of human mesenchymal stem cells. Stem Cells. 2014;32(6):1408–1419.

48. Zhang LZ, Huang LY, Huang AL, Liu JX, Yang F. CRIP1 promotes cell migration, invasion and epithelial-mesenchymal transition of cervical cancer by activating the Wnt/beta□catenin signaling pathway. Life Sci. 2018;207:420–427.

49. Fevr T, Robine S, Louvard D, Huelsken J. Wnt/beta-catenin is essential for intestinal homeostasis and maintenance of intestinal stem cells. Mol Cell Biol. 2007;27(21):7551–7559.

50. Nusse R. Wnt signaling and stem cell control. Cell Res. 2008;18(5):523–527.

51. Mas A, Cervello I, Gil-Sanchis C, Faus A, Ferro J, Pellicer A, Simon C. Identification and characterization of the human leiomyoma side population as putative tumor-initiating cells. Fertil Steril. 2012;98(3):741–751 e746.

52. Paul EN, Burns GW, Carpenter TJ, Grey JA, Fazleabas AT, Teixeira JM. Transcriptome Analyses of Myometrium from Fibroid Patients Reveals Phenotypic Differences Compared to Non-Diseased Myometrium. Int J Mol Sci. 2021;22(7).

53. Tang L, Bergevoet SM, Gilissen C, de Witte T, Jansen JH, van der Reijden BA, Raymakers RA. Hematopoietic stem cells exhibit a specific ABC transporter gene expression profile clearly distinct from other stem cells. BMC Pharmacol. 2010;10:12.

54. Bhartiya D. Adult tissue-resident stem cells-fact or fiction? Stem Cell Res Ther. 2021;12(1):73.

55. Zhou X, Lindsay H, Robinson MD. Robustly detecting differential expression in RNA sequencing data using observation weights. Nucleic Acids Res. 2014;42(11):e91.

56. Hao Y, Hao S, Andersen-Nissen E, Mauck WM, 3rd, Zheng S, Butler A, Lee MJ, Wilk AJ, Darby C, Zager M, Hoffman P, Stoeckius M, Papalexi E, Mimitou EP, Jain J, Srivastava A, Stuart T, Fleming LM, Yeung B, Rogers AJ, McElrath JM, Blish CA, Gottardo R, Smibert P, Satija R. Integrated analysis of multimodal single-cell data. Cell. 2021;184(13):3573–3587 e3529.

57. Arora R, Fries A, Oelerich K, Marchuk K, Sabeur K, Giudice LC, Laird DJ. Insights from imaging the implanting embryo and the uterine environment in three dimensions. Development. 2016;143(24):4749–4754.

58. Flores D, Madhavan M, Wright S, Arora R. Mechanical and signaling mechanisms that guide pre-implantation embryo movement. Development. 2020;147(24).

59. Benjamini Y, Hochberg Y. Controlling the False Discovery Rate - a Practical and Powerful Approach to Multiple Testing. J Roy Stat Soc B Met. 1995;57(1):289–300.

