## Supplementary figures and images for "Cysteine-Rich Intestinal Protein 1 is a Novel Surface Marker for Myometrial Stem/Progenitor Cells"

### Fig S1

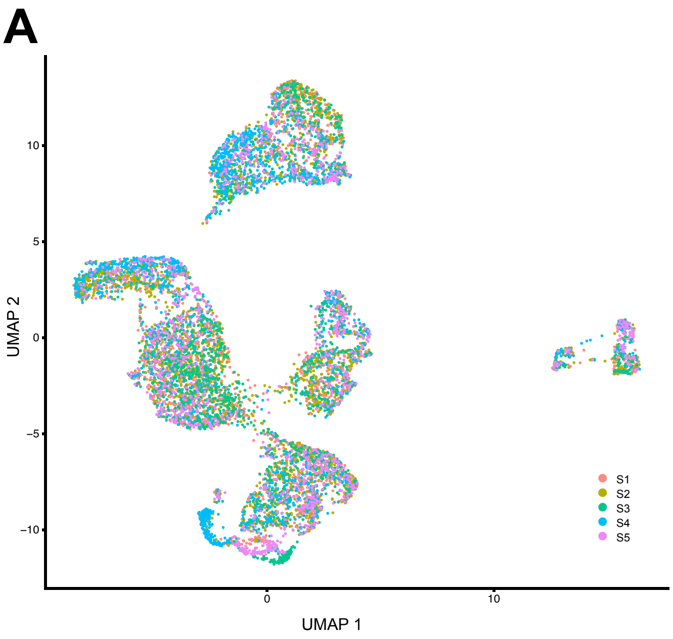

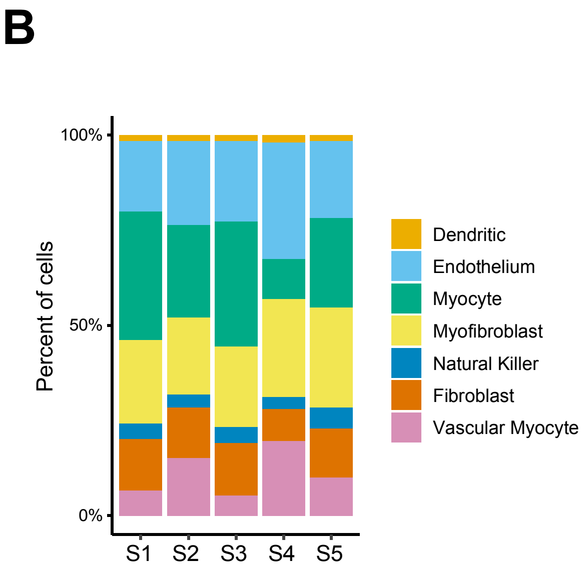

### Fig S2

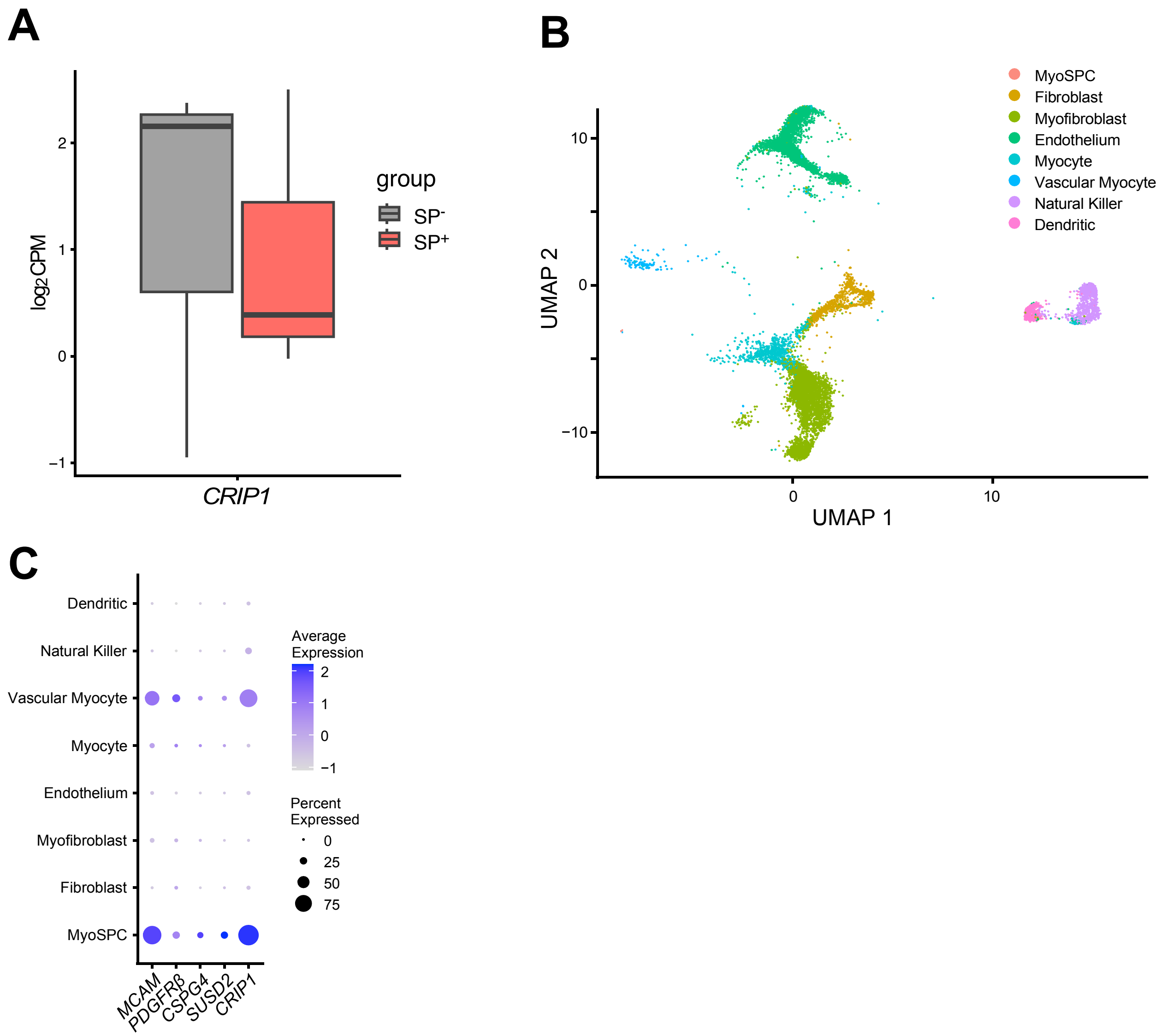
